# The amoeboid migration of monocytes in confining channels requires the local remodeling of the cortical actin cytoskeleton by cofilin-1

**DOI:** 10.1101/2023.08.11.553020

**Authors:** Maria F. Ullo, Anna E. D’Amico, Sandrine B. Lavenus, Jeremy S. Logue

## Abstract

Within the bloodstream, monocytes must traverse the microvasculature to prevent leukostasis, which is the entrapment of monocytes within the confines of the microvasculature. Using the model cell line, THP-1, and VCAM-1 coated channels to simulate the microvasculature surface, we demonstrate that monocytes predominantly adopt an amoeboid phenotype, which is characterized by the formation of blebs. As opposed to cortical actin flow in leader blebs, cell movement is correlated with myosin contraction at the cell rear. It was previously documented that cofilin-1 promotes cortical actin turnover at leader bleb necks in melanoma cells. In monocytes, our data suggest that cofilin-1 promotes the local upregulation of myosin contractility through actin cytoskeleton remodeling. In support of this concept, cofilin-1 is found to localize to a single cell edge. Moreover, the widespread upregulation of myosin contractility was found to inhibit migration. Thus, monocytes within the microvasculature may avoid entrapment by adopting an amoeboid mode of migration.

**Summary Statement:** In confining channels, monocytes largely adopt an amoeboid migrating phenotype, which is found to depend on the upregulation of myosin contractility at the cell rear and cortical actin remodeling by cofilin-1.

## Introduction

In response to inflammatory cues (*e.g*., chemokines), monocytes extravasate and migrate towards a site of infection or tissue injury where they differentiate into macrophages or dendritic cells^1-4^. To survey tissues, however, monocytes must freely circulate in the bloodstream. The lumen of a micro-vessel is very small, ranging from 4 to 9 μm in diameter^5^. Accordingly, monocytes within microvasculature are challenged with moving through a physically confining environment^6^. The entrapment of monocytes within the microvasculature, termed leukostasis, results in respiratory and neurological distress^7^. How monocytes avoid entrapment within the microvasculature, however, is not well understood.

Cells can migrate in varied ways. This includes mesenchymal migration, which is characterized by actin polymerization-based protrusion of the leading edge and integrin adhesion to the extracellular matrix (ECM)^8^. During amoeboid migration, protrusion of the leading edge is driven by intracellular pressure and cytoplasmic flow^9^. In cancer cells, for instance, high intracellular pressure and cytoplasmic flow drive the formation of plasma membrane (PM) blebs^10-12^. Under 2 dimensional (2D) or vertical confinement, cancer cells will often adopt fast amoeboid migration^10^. During fast amoeboid migration, the motive force for cell movement is generated by leader blebs, which contain a rapid cortical actin flow^12^. In melanoma cells, flow depends on a high rate of cortical actin turnover at bleb necks. We previously found that the actin severing factors, ADF and cofilin-1, are required for a high rate of cortical actin flow in leader blebs^13^. In some cells, blebs had protracted necks and failed to retract; thus, ADF and cofilin-1 were shown to be essential for normal bleb dynamics^13, 14^. Importantly, fast amoeboid migration only requires friction between the cell and microenvironment for force transmission^15^. In heterogenous tissues, cells may use fast amoeboid migration for passing between highly and poorly adhesive environments.

Without chemokines, integrin levels in monocytes are very low and they survive in suspension (*i.e*., without substrate adhesion)^16^. To traverse the microvasculature and avoid entrapment, we wondered if monocytes may adopt an integrin-independent mode of migration. Previous studies have shown that immune cells can migrate within interstitial space in the absence of integrins^17, 18^. Using the acute monocytic leukemia cell line, THP-1, as a model we demonstrate that monocytes adopt an amoeboid mode of migration in confining channels coated with VCAM-1. Strikingly, cell movement is found to correlate with a high level of myosin contractility at the cell rear. In monocytes, our data suggest that cofilin-1 promotes the local upregulation of myosin contractility. Thus, the mechanism by which cofilin-1 promotes amoeboid migration in monocytes is distinct from melanoma cells and may help monocytes avoid entrapment within the microvasculature.

## Results

In the absence of inflammatory cues (*e.g*., chemokines), monocytes have very low levels of integrins and do not adhere to substrates^16^. To determine if monocytes may use an integrin-independent, *i.e*., amoeboid, mode of migration we first quantified the percent of cells that display blebs in suspension culture, as plasma membrane (PM) blebbing is a hallmark of amoeboid migration^19^. Using high-resolution imaging, ∼25% of THP-1 monocyte-like cells are observed blebbing in suspension culture (Fig. 1A). Previously, the 2 dimensional (2D) or vertical confinement of cancer cells was shown to trigger a phenotypic transition to fast amoeboid migration in cancer cells^10-12^. Fast amoeboid migration is driven by a leader bleb, which contains a rapid cortical actin flow^12^. Thus, we wondered if monocytes may undergo a similar phenotypic transition. Although confinement down to 3 μm by a polydimethylsiloxane (PDMS) ceiling potently increased the percentage of blebbing cells, THP-1 monocyte-like cells could not be seen forming leader blebs (Fig. 1A’). Within tissues, monocytes are challenged with traversing the confines of the microvasculature. Accordingly, we subjected monocytes to confinement in channels. Microchannels are 3, 8, and 100 μm in height, width, and length (HWL), respectively, and coated with VCAM-1 (1 μg/mL). To visualize filamentous-actin (F-actin), we transduced THP-1 monocyte-like cells with a lentivirus encoding an enhanced green fluorescent protein (EGFP) tagged version of LifeAct^20^. In microchannels, we observed numerous small blebs primarily at the leading edge of motile monocytes (Fig. 1B-C & Movie 1). Strikingly, we observed that cell movement correlated with increased curvature at the cell rear, which is a shape change suggestive of a high level of myosin contractility (Fig. 1B-D & Movie 1). In agreement with this concept, the level of an EGFP tagged version of the regulatory light chain (RLC) of myosin was found to be high at the trailing edge of a motile cell (Fig. 1E-H & Movie 2). Conversely, poorly motile cells (*i.e*., moving less than half of the cell body length over 5 hr) did not display increased curvature at a single cell edge (Movie 3). Thus, the migration of monocytes correlates with the local upregulation of myosin at the cell rear and not cortical actin flow in leader blebs.

**Figure 1.**
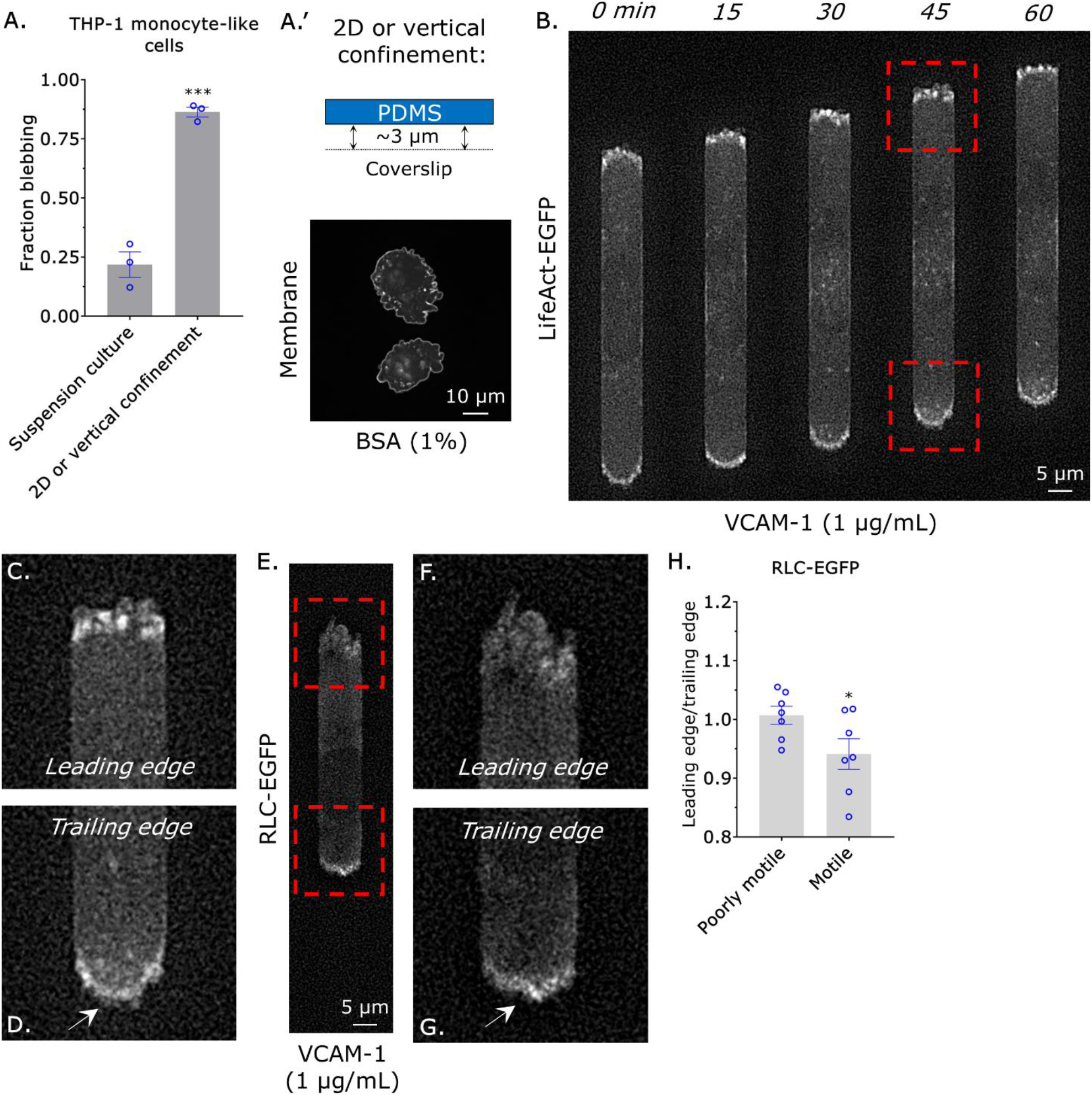
Monocyte migration correlates with contraction at the cell rear. **A**. Fraction of cells displaying blebs in suspension culture or when confined down to 3 μm by a PDMS ceiling, as measured by live imaging of THP-1 monocyte-like cells stained with a membrane dye. Statistical significance was determined by a t-test (mean +/- SEM). A representative image of cells confined down to 3 μm by a PDMS ceiling stained with membrane dye is shown in (A’). For 2D or vertical confinement, PDMS is coated with bovine serum albumin (BSA; 1%). **B**. Montage of a THP-1 monocyte-like cell with the F-actin marker, LifeAct-EGFP, migrating within a confining channel. **C**. Zoom from (B; *top*) showing multiple small blebs at the leading edge. **D**. Zoom from (B; *bottom*) showing contraction of the cell rear, as indicated by increased curvature. **E**. Localization of myosin in a motile cell, as indicated by RLC-EGFP. **F**. Zoom from (E; *top*) showing a low level of RLC-EGFP at the leading edge. **G**. Zoom from (E; *bottom*) showing a high level of RLC-EGFP at the trailing edge. **H**. Ratio of RLC-EGFP at the leading edge over the trailing edge in poorly motile (*i.e*., moving less than half of the cell body length over 5 hr) and motile cells. Statistical significance was determined by a t-test (mean +/- SEM). Microchannels are coated with VCAM-1 (1 μg/mL) and are 3 μm in height, 8 μm in width, and 100 μm in length. * - p ≤ 0.05, ** - p ≤ 0.01, *** - p ≤ 0.001, and **** - p ≤ 0.0001

Previously, we documented that the actin severing factors, ADF and cofilin-1, are essential for cortical actin flow in leader blebs^13^. As motile monocytes form numerous blebs at the leading edge, we wondered what role they may have in regulating the amoeboid migration of monocytes. By RT-qPCR, we found that THP-1 monocyte-like cells predominantly contain cofilin-1 (Fig. 2A). We then determined to what degree cofilin-1 regulates the total level of F-actin. To measure the total level of F-actin, monocytes in suspension were paraformaldehyde (PFA) treated and stained with fluorescently conjugated phalloidin and subjected to flow cytometry analysis. This method is preferred, as preserving the actin cytoskeleton with PFA in many cells within microchannels with high fidelity is technically challenging. As validation, we treated monocytes with the actin depolymerizing drug, Latrunculin (5 μM), before flow cytometry^21^. Compared to untreated, F-actin levels were reduced by ∼95% in Latrunculin treated cells (Fig. 2B). While removing ADF by RNA interference (RNAi) had no effect, removing cofilin-1 led to a ∼10% increase in the total level of F-actin (Fig. 2B-C). Moreover, RNAi of cofilin-1 and ADF did not have an additive effect (Fig. 2B-C). As cofilin-1 was found to regulate the total level of F-actin in monocytes, we wondered how cofilin-1 might regulate amoeboid migration. Qualitatively, bleb dynamics were unchanged after cofilin-1 RNAi (Fig. 2D-F & Movie 4-5). Upon closer inspection, monocytes failed to form a contractile edge after cofilin-1 RNAi, which was found to correlate with migration (Fig. 2D-F & Movie 4-5). Cell tracking revealed a ∼50% decrease in instantaneous speed after cofilin-1 RNAi (Fig. 2G). Similarly, the directionality ratio over elapsed time of cells after cofilin-1 RNAi was significantly reduced (Fig. 2H). Therefore, cofilin-1 may regulate myosin contractility through actin cytoskeleton remodeling in monocytes.

**Figure 2.**
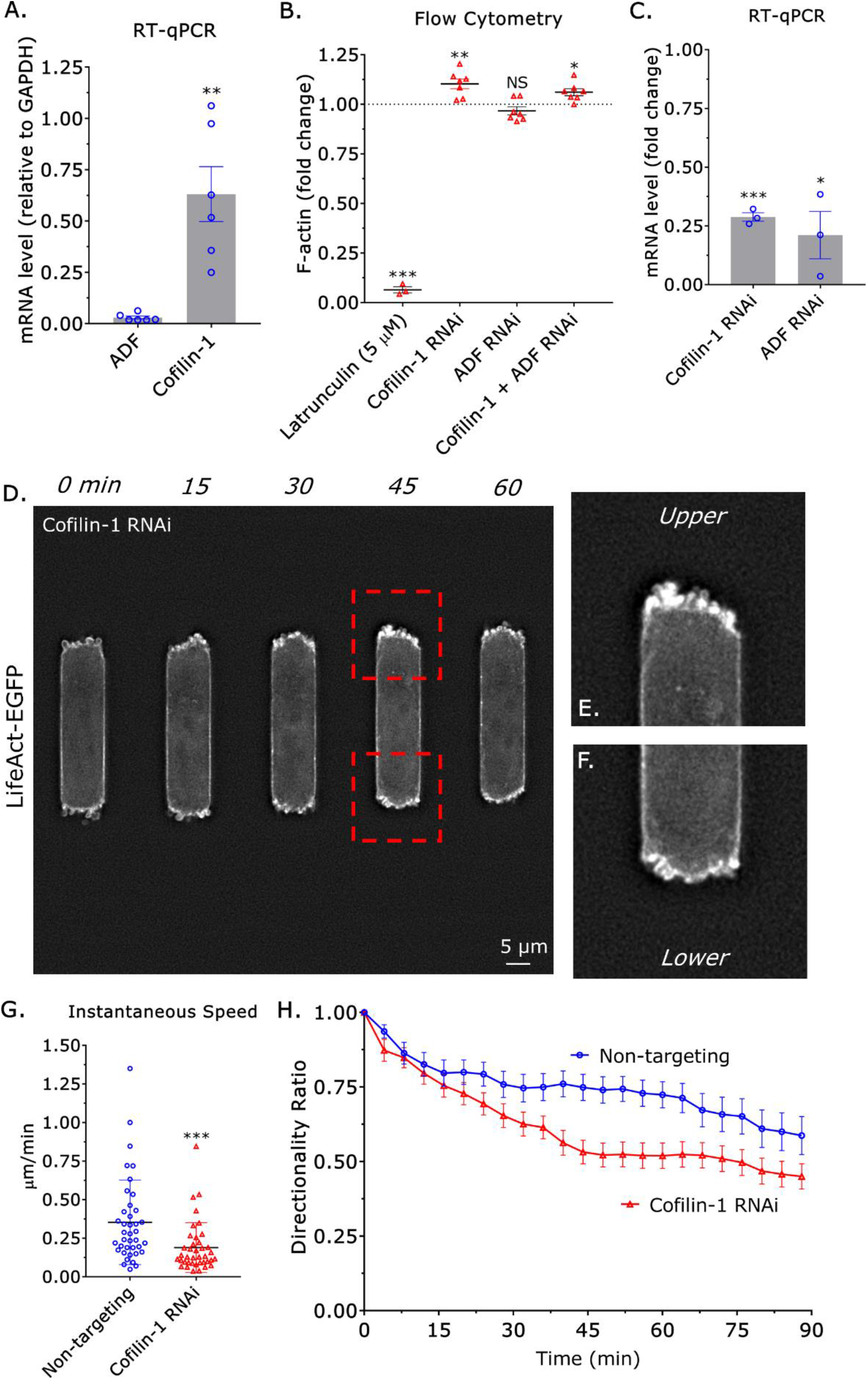
Cofilin-1 promotes the amoeboid migration of monocytes in confining channels. **A**. As measured by RT-qPCR, THP-1 monocyte-like cells predominantly contain cofilin-1 and not ADF. Statistical significance was determined by a t-test (mean +/- SEM). **B**. The total level of F-actin for latrunculin (5 μM; positive control), cofilin-1, ADF, and cofilin-1 + ADF siRNA treated cells, as measured by flow cytometry of phalloidin stained cells. Statistical significance was determined by a one-sample t-test (theoretical value = 1) (mean +/- SEM). **C**. Confirmation of cofilin-1 and ADF RNAi by RT-qPCR. Statistical significance was determined by a one-sample t-test (theoretical value = 1) (mean +/- SEM). **D**. Montage of a THP-1 monocyte-like cell with the marker of F-actin, LifeAct-EGFP, and treated with a cofilin-1 siRNA. **E**. Zoom from (D; *top*) showing multiple small blebs and little contraction, as indicated by the lack of curvature at the cell edge. **F**. Zoom from (D; *bottom*) showing multiple small blebs and little contraction, as indicated by the lack of curvature at the cell edge. **G**. Instantaneous speeds for non-targeting and cofilin-1 siRNA treated cells, as measured by manual tracking. Statistical significance was determined by a t-test (mean +/- SD). **H**. Directionality ratio over elapsed time for non-targeting and cofilin-1 siRNA treated cells, as measured by manual tracking (mean +/- SEM). Microchannels are coated with VCAM-1 (1 μg/mL) and are 3 μm in height, 8 μm in width, and 100 μm in length. * - p ≤ 0.05, ** - p ≤ 0.01, *** - p ≤ 0.001, and **** - p ≤ 0.0001.

In cells, cofilin-1 is heavily regulated by the Ser/Thr kinases, LIM Kinase (LIMK) 1 and 2^22^. Phosphorylation by either kinase on Ser 3 of cofilin-1 leads to its inactivation^22^. Therefore, we determined if removing LIMK1 or 2 by RNAi had any effect on monocyte migration in confining channels. By RT-qPCR, we found that THP-1 monocyte-like cells predominantly contain LIMK1 (Fig. 3A). Surprisingly, while there is a slight tendency for cells after LIMK1 RNAi to have a higher instantaneous speed and directionality ratio over elapsed time, these differences are not statistically significant (Fig. 3B-D). Accordingly, cofilin-1 may not be subject to strong negative regulation by LIMK1 in monocytes.

**Figure 3.**
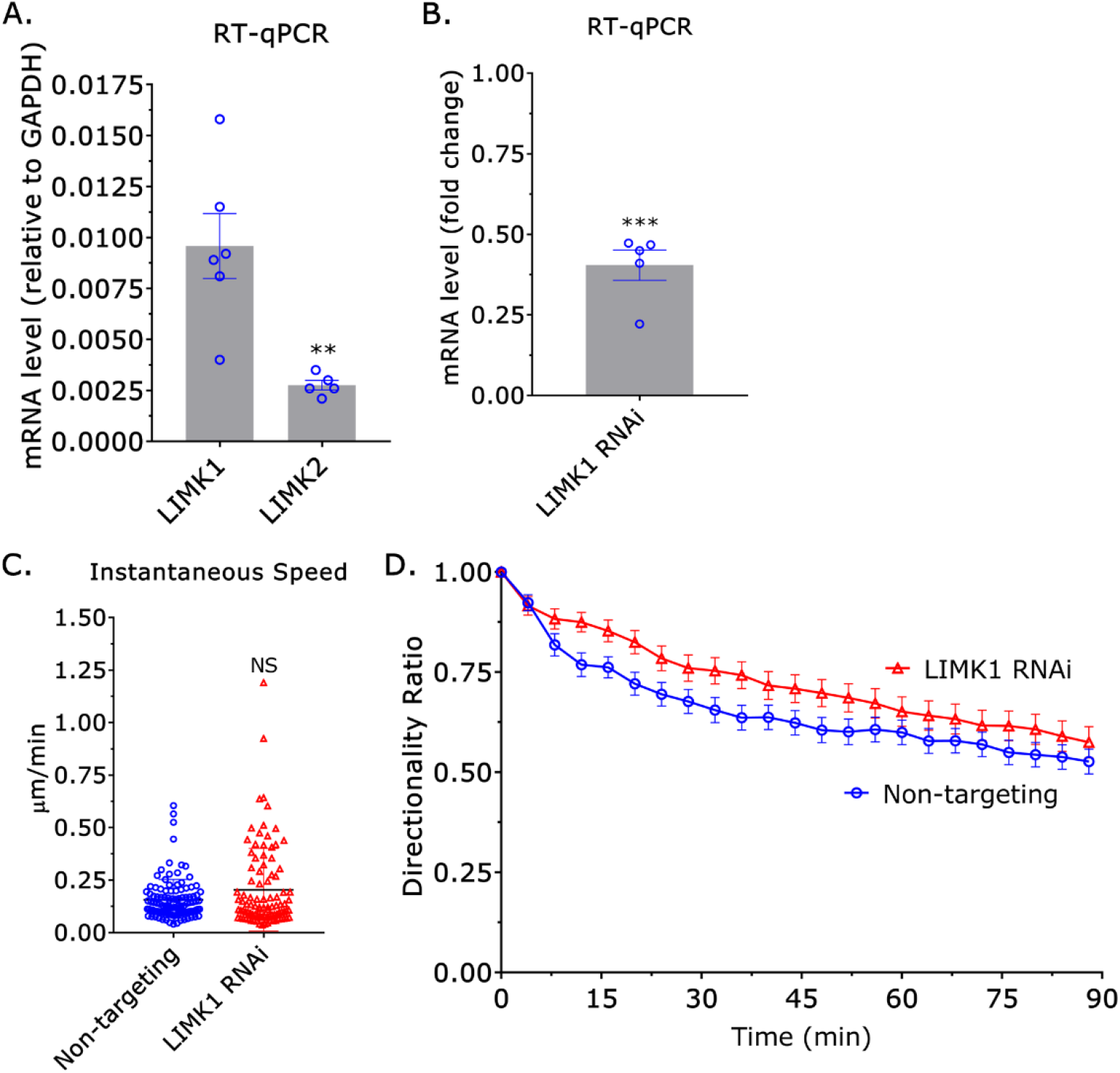
Cofilin-1 is not heavily regulated by LIMK1. **A**. By RT-qPCR, THP-1 monocyte-like cells predominantly contain LIMK1 and not LIMK2. Statistical significance was determined by a t-test (mean +/- SEM). **B**. Confirmation of LIMK1 RNAi by RT-qPCR. Statistical significance was determined by a one-sample t-test (theoretical value = 1) (mean +/- SEM). **C**. Instantaneous speeds for non-targeting and cofilin-1 siRNA treated cells, as measured by manual tracking. Statistical significance was determined by a t-test (mean +/- SD). **D**. Directionality ratio over elapsed time for non-targeting and LIMK1 siRNA treated cells, as measured by manual tracking (mean +/- SEM). Microchannels are coated with VCAM-1 (1 μg/mL) and are 3 μm in height, 8 μm in width, and 100 μm in length. * - p ≤ 0.05, ** - p ≤ 0.01, *** - p ≤ 0.001, and **** - p ≤ 0.0001

As migration correlated with the local upregulation of myosin contractility at the cell rear, we wondered if actin remodeling by cofilin-1 could promote myosin contractility. To measure the stiffness of cells, we used a previously described approach that involves measuring the height (*h*) and diameter (*d*) of cells sandwiched between two polyacrylamide (PA) gels of known stiffness (1 kPa)^10^. While removing ADF by RNAi had no effect, removing cofilin-1 led to a ∼25% decrease in stiffness (*h*/*d*; Fig. 4A). Moreover, removing cofilin-1 and ADF did not have an additive effect (Fig. 4A). Thus, our data suggest that cofilin-1 promotes migration through the upregulation of myosin contractility. To ascertain if the widespread upregulation of myosin contractility promotes migration, we removed myosin phosphatase target subunit 1 (MYPT1) by RNAi. In the absence of MYPT1, the regulatory light chain (RLC) of myosin remains phosphorylated, increasing contractility^10^. Surprisingly, the widespread upregulation of myosin contractility by MYPT1 RNAi led to a ∼50% decrease in instantaneous speed (Fig. 4B & D). Similarly, the directionality ratio over elapsed time of cells after MYPT1 RNAi was significantly reduced (Fig. 4C-D). As opposed to widespread upregulation, cofilin-1 may promote migration through the local upregulation of myosin contractility. Therefore, we determined if cofilin-1 is localized to a single cell edge. In motile cells, cofilin-1-EGFP was mostly cytosolic; however, pseudo color representation revealed an enrichment of cofilin-1-EGFP at the trailing edge (Fig. 4E-F & Movie 6). To bolster these results, we localized endogenous cofilin-1 in monocytes loosely attached to poly-L-lysine coated cover glass by immunofluorescence (IF). Strikingly, endogenous cofilin-1 was found to consistently localize to a single edge (Fig. 4G-H). Interestingly, the total level of F-actin at sites of cofilin-1 enrichment was reduced, possibly because of increased turnover (Fig. 4G-H)^23^. Therefore, our data suggest that cofilin-1 locally regulates the actin cytoskeleton in monocytes, which promotes the formation of a contractile cell rear and migration.

**Figure 4.**
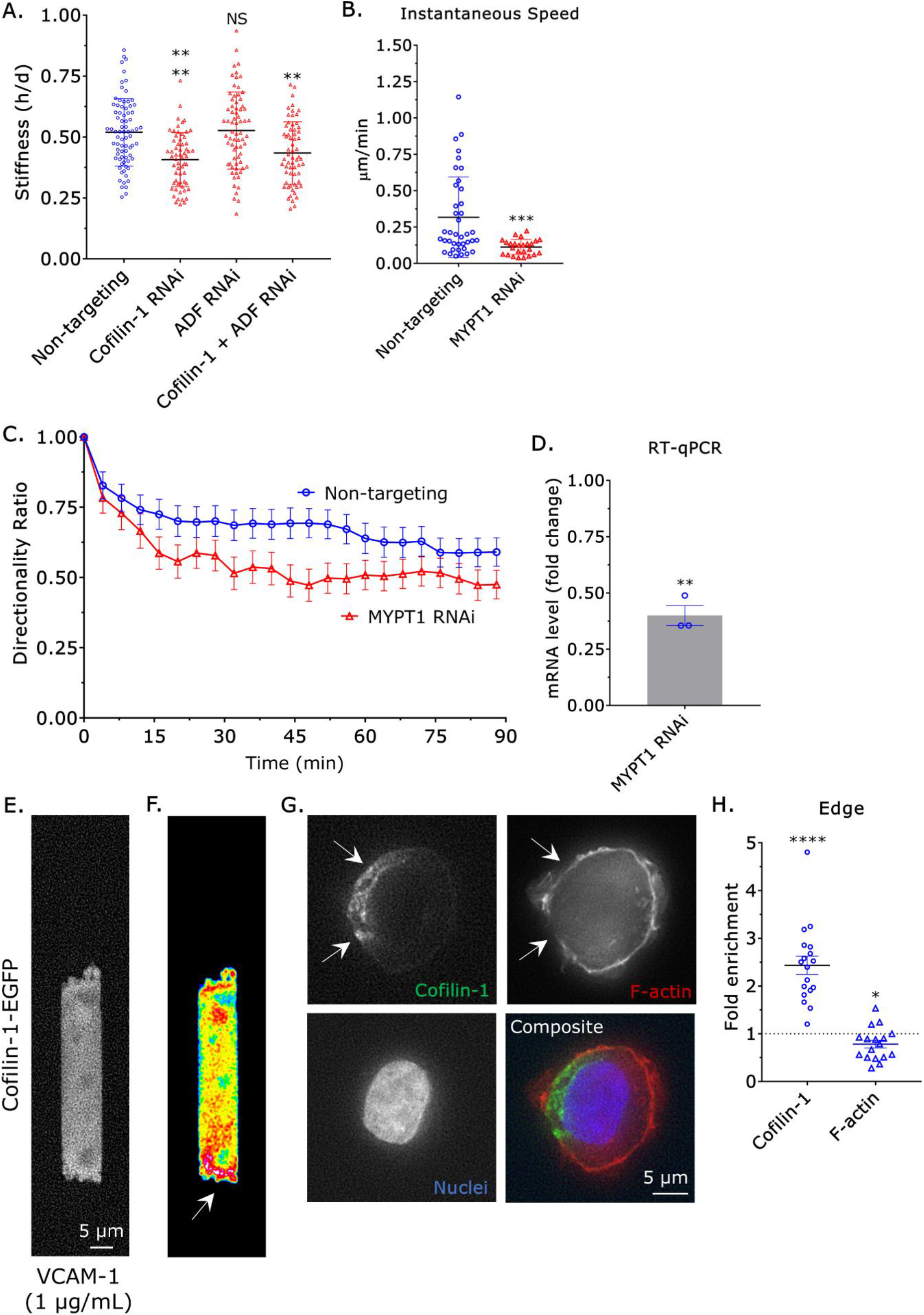
Widespread up-regulation of myosin contractility inhibits migration. **A**. Cell stiffnesses for non-targeting, cofilin-1, ADF, and cofilin-1 + ADF siRNA treated cells, as measured by a gel sandwich assay. Statistical significance was determined by a multiple comparisons test (mean +/- SD). **B**. Instantaneous speeds for non-targeting and MYPT1 siRNA treated cells, as measured by manual tracking. Statistical significance was determined by a t-test (mean +/- SD). **C**. Directionality ratio over elapsed time for non-targeting and MYPT1 siRNA treated cells, as measured by manual tracking (mean +/- SEM). **D**. Confirmation of MYPT1 RNAi by RT-qPCR. Statistical significance was determined by a one-sample t-test (theoretical value = 1) (mean +/- SEM). **E**. Localization of cofilin-1 in a motile cell, as indicated by cofilin-1-EGFP. **F**. Pseudo color image of (E), which highlights the high level of cofilin-1-EGFP at the trailing edge. **G**. Immunofluorescence (IF) of endogenous cofilin-1 in a THP-1 monocyte-like cell loosely adhered to a poly-L-lysine coated coverslip. F-actin staining appears weaker in regions where cofilin-1 is enriched (*arrows*). **H**. Quantitative evaluation of cofilin-1 and F-actin fold enrichment at cell edges. Statistical significance was determined by a one-sample t-test (theoretical value = 1) (mean +/- SEM). Microchannels are coated with VCAM-1 (1 μg/mL) and are 3 μm in height, 8 μm in width, and 100 μm in length. * - p ≤ 0.05, ** - p ≤ 0.01, *** - p ≤ 0.001, and **** - p ≤ 0.0001

In response to inflammatory cues (*e.g*., chemokines), monocytes extravasate and migrate to sites of infection or tissue injury where they differentiate into macrophages or dendritic cells. Using RT-qPCR, we determined if actin turnover or polymerization factors are up- or downregulated after differentiation into THP-1 macrophage-like cells. Interestingly, after differentiation the level of the Ca^2+^ regulated actin turnover factor, gelsolin, increased by ∼5-fold (Fig. 5A)^22^. While ADF levels were unchanged, cofilin-1 levels increased by ∼2-fold after differentiation (Fig. 5A). The levels of the cofilin-1 regulators, LIMK1, LIMK2, and SSH1 were unchanged after differentiation (Fig. 5B). We then measured the mRNA levels of Arp2/3 and several formins before and after differentiation. Consistent with the assumption of a phenotype with wide lamellipodia, which THP-1 macrophage-like cells frequently display, Arp2 and 3 were found to be up-regulated by ∼3-fold after differentiation (Fig. 5C). Interestingly, formin like 1 (FMNL1) was found to be up-regulated by ∼3-fold, whereas formin like 2 (FMNL2) was dramatically up-regulated (Fig. 5C). Based on these data, cofilin-1, gelsolin, Arp2/3, and FMNL1/2 may have essential function(s) in THP-1 macrophage-like cells.

**Figure 5.**
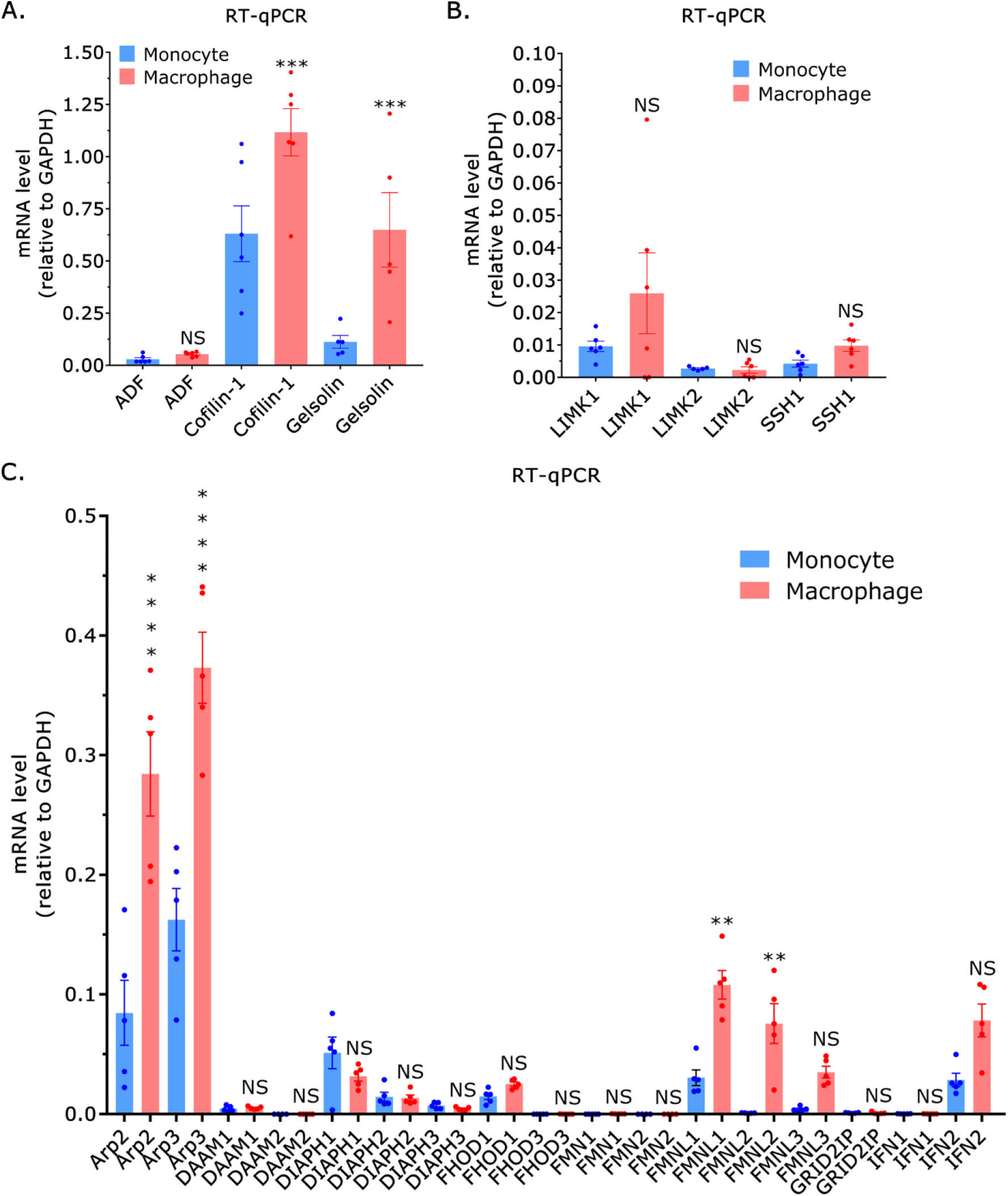
THP-1 macrophage-like cells possess a unique set of actin cytoskeletal regulators. **A**. By RT-qPCR, THP-1 macrophage-like cells have more of the actin turnover factors, cofilin-1, and gelsolin. Statistical significance was determined by a multiple comparisons test (mean +/- SEM). **B**. By RT-qPCR, the levels of LIMK1, LIMK2, and SSH1 are unchanged after differentiation. Statistical significance was determined by a multiple comparisons test (mean +/- SEM). **C**. By RT-qPCR, THP-1 macrophages have more of the actin polymerization factors, Arp2/3, and FMNL1/2. Statistical significance was determined by a multiple comparisons test (mean +/- SEM). * - p ≤ 0.05, ** - p ≤ 0.01, *** - p ≤ 0.001, and **** - p ≤ 0.0001

## Discussion

To survey tissues, monocytes must traverse the confines of the microvasculature. As monocytes survive in suspension (*i.e*., without substrate adhesion), we wondered if monocytes could use amoeboid migration to avoid entrapment within the microvasculature. Within confining channels, we found that the model cell line, THP-1, almost entirely adopted an amoeboid phenotype. Previously, we found that melanoma cells frequently adopted a hybrid phenotype in channels coated with fibronectin (10 μg/mL)^24^. Cells displaying the hybrid phenotype had hallmarks of mesenchymal (*e.g*., focal adhesions) and amoeboid (*e.g*., blebs) migration^8^. The absence of mesenchymal features in monocytes, however, is consistent with their low level of substrate adhesion^16^.

As migration correlated with contraction of the cell rear, we determined what actin cytoskeletal remodeling factors may promote motility. Previously, we demonstrated that the actin severing factors, ADF and cofilin-1, are required for the rapid cortical actin flow in leader blebs^13^. Cancer cells frequently form leader blebs under 2D or vertical confinement, which drives fast amoeboid migration^10-12^. Therefore, we wondered if ADF or cofilin-1 have a similar role in amoeboid migrating monocytes. Like what we previously reported in melanoma cells, RNAi of cofilin-1 but not ADF increased the total level of F-actin^13^. Moreover, removing cofilin-1 and ADF did not have an additive effect. Despite monocytes not using leader blebs for fast amoeboid migration, RNAi of cofilin-1 led to a significant decrease in instantaneous speeds and directionality. These data, therefore, suggest that there is a distinct mechanism by which cofilin-1 promotes the amoeboid migration of monocytes.

Through its actin cytoskeletal remodeling activity, cofilin-1 has been shown to promote myosin contractility^25, 26^. Therefore, we determined if cofilin-1 may promote myosin contractility in monocytes. Indeed, we found that RNAi of cofilin-1 but not ADF led to a decrease in cell stiffness. Moreover, removing cofilin-1 and ADF did not have an additive effect. We then wondered if the widespread upregulation of myosin contractility could promote the amoeboid migration of monocytes. To accomplish this, we depleted cells of MYPT1 by RNAi. Strikingly, removing MYPT1 led to a significant decrease in instantaneous speeds and directionality. These data suggest that migration requires a tension gradient created by a local increase in myosin contractility. Similarly, the establishment of the cell rear is reinforced by myosin in neutrophils, whereas dendritic cells require a high level of myosin contractility at the cell rear for migration specifically through confined environments^27-29^. As migration correlated with the contraction of the cell rear, cofilin-1 may promote migration by locally upregulating myosin contractility. Although cofilin-1-EGFP was largely cytosolic in migrating monocytes, we were able to consistently observe endogenous cofilin-1 localizing to a single cell edge by IF. Thus, cofilin-1 may promote the amoeboid migration of monocytes through the local upregulation of myosin contractility.

In response to inflammatory cues (*e.g*., chemokines), monocytes extravasate and migrate to sites of infection or injury where they differentiate into macrophages or dendritic cells. Therefore, macrophages or dendritic cells are likely to have a unique set of actin cytoskeletal regulators. Interestingly, gelsolin was upregulated by ∼5-fold after differentiation into THP-1 macrophage-like cells. Gelsolin is a Ca^2+^ regulated actin turnover factor; thus, the actin cytoskeleton in macrophages may be poised to respond to intracellular Ca^2+^ signaling^30^. Arp2/3 and FMNL1/2 were also found to be upregulated in THP-1 macrophage-like cells. Arp2/3 is an important regulator of actin dynamics during phagocytosis^31^. Notably, Arp2/3 activity has been found to be dispensable for confined migration in melanoma cells^24, 32^. FMNL1/2 have been shown to promote the formation of podosomes in macrophages^33^. On the other hand, high levels of cofilin-1 mRNA could be found in macrophages and monocytes. Although cofilin-1 is likely to play differential roles in monocytes and macrophages, this work further demonstrates the pivotal role(s) that cofilin-1 has in regulating the actin cytoskeleton in diverse cell types.

## Experimental Procedures

### Cell culture

THP-1 (TIB-202) were obtained from the American Type Culture Collection (ATCC; Manassas, VA). Cells were cultured in RPMI (Thermo Fisher, Carlsbad, CA) supplemented with fetal bovine serum (FBS; 10%) (cat no. 12106C; Sigma Aldrich, St. Louis, MO), L-glutamine (cat no. 35050061; Thermo Fisher), antibiotic-antimycotic (cat no. 15240096; Thermo Fisher), and β-mercaptoethanol (50 μM).

### Plasmids

Plasmids encoding LifeAct-EGFP (cat no. 51010), RLC-EGFP (cat no. 35680), and cofilin-1-EGFP (cat no. 50859) were purchased from Addgene (Watertown, MA).

### RT-qPCR

Total RNA was isolated from cells using the PureLink RNA Mini Kit (cat no. 12183018A; Thermo Fisher) and was used for reverse transcription using a High-Capacity cDNA Reverse Transcription Kit (cat no. 4368814; Thermo Fisher). qPCR was performed using PowerUp SYBR Green Master Mix (cat no. A25742; Thermo Fisher) on a real-time PCR detection system (CFX96; Bio-Rad, Hercules, CA). Relative mRNA levels were calculated using the ΔCt method. All qPCR primer pair sequences were from OriGene (Rockville, MD).

### RNA interference (RNAi)

Non-targeting (cat no. 4390844), ADF (cat no. 4392422; s21737), cofilin-1 (cat no. 4392420; s2936), LIMK1 (cat no. 4390826; s8188), and MYPT1 (cat no. 4390824; s9235) siRNAs were purchased from Thermo Fisher. All siRNA transfections were performed using RNAiMAX (5 μL; Thermo Fisher) and OptiMEM (400 μL; Thermo Fisher). 250,000 cells were collected and seeded in 6-well plates in complete media. siRNAs in RNAiMAX/OptiMEM were added to cells in complete media (2 mL) at a final concentration of 50 nM. Cells were incubated with siRNAs for 3 days.

### Flow cytometry

1 × 10^6^ cells in HBS were fixed using 4% paraformaldehyde (PFA) (cat no. 15710; Electron Microscopy Sciences, Hatfield, PA) for 20 min at room temperature. After washing, cell pellets were resuspended in flow buffer (HBS with 1% BSA) and incubated with 0.1% Triton X-100, a 1:250 dilution of Alexa Fluor 568-conjugated phalloidin (cat no. A22287; Thermo Fisher), and 1:1000 dilution of DAPI (cat no. D1306; Thermo Fisher) for 1 hr at room temperature. Cells were then re-pelleted, washed, and suspended in flow buffer. Data were acquired on a FACSCalibur (BD Biosciences, Franklin Lakes, NJ) flow cytometer. Flow cytometric analysis was performed using FlowJo (Ashland, OR) software.

### 2D confinement

This protocol has been described in detail elsewhere^34^. Briefly, PDMS (cat no. 24236-10) was purchased from Electron Microscopy Sciences (Hatfield, PA). 2 mL was cured overnight at 37 °C in each well of a 6-well glass bottom plate (cat no. P06-1.5H-N; Cellvis, Mountain View, CA). Using a biopsy punch (cat no. 504535; World Precision Instruments, Sarasota, FL), an 8 mm hole was cut and 3 mL of serum free media containing 1% BSA was added to each well and incubated overnight at 37 °C. After removing the serum free media containing 1% BSA, 300 μL of complete media containing trypsinized cells (250,000 to 1 million) and 2 μL of 3.11 μm beads (cat no. PS05002; Bangs Laboratories, Fishers, IN) were then pipetted into the round opening.

The vacuum created by briefly lifting one side of the hole with a 1 mL pipette tip was used to move cells and beads underneath the PDMS. Finally, 3 mL of complete media was added to each well and cells were recovered for ∼60 min before imaging.

### Microchannel preparation

PDMS (cat no. 24236-10; Electron Microscopy Sciences) was prepared using a 1:7 ratio of base and curing agent. Uncured PDMS was poured over the wafer mold, placed in a vacuum chamber to remove bubbles, moved to a 37 °C incubator, and left to cure overnight. After curing, small PDMS slabs with microchannels were cut using a scalpel, whereas cell loading ports were cut using a 0.4 cm hole punch (cat no. 12-460-409; Fisher Scientific, Hampton, NH).

For making PDMS coated cover glass (cat no. 12-545-81; Fisher Scientific), 30 μL of uncured PDMS was pipetted at the center of the cover glass, placed in a modified mini-centrifuge, and spun for 30 sec for even spreading. The PDMS coated cover glass was then cured for at least 1 hr on a 95 °C hot plate.

Prior to slab and coated cover glass joining, PDMS surfaces were activated for ∼1 min by plasma treatment (cat no. PDC-32G; Harrick Plasma, Ithaca, NY). Immediately after activation, slabs were bonded to coated cover glass. For complete bonding, the apparatus was incubated at 37 °C for at least 1 hr.

### Microchannel coating

Immediately after being plasma treated for ∼ 1 min, VCAM-1 (cat no. 862-VC; R&D Systems, Minneapolis, MN) dissolved in PBS at a concentration of 1 μg/mL was pumped into microchannels using a modified motorized pipette. To remove any bubbles pumped into microchannels, the apparatus was left to coat in a vacuum chamber for at least 1 hr. Afterward, VCAM-1 solution was aspirated out and microchannels were rinsed twice by pumping in PBS. Finally, microchannels were incubated in complete media overnight at 4 °C before use.

### Microchannel loading

Prior to cells being loaded into microchannels, complete media was aspirated, and microchannels were placed into an interchangeable cover-glass dish (cat no. 190310-35; Bioptechs, Butler, PA). Cells in 300 μL of complete media, stained with 1 μL of far-red membrane dye (cat no. C10046; Thermo Fisher), were pumped into microchannels using a modified motorized pipette. Once at least 20 cells are observed in microchannels by low magnification brightfield microscopy, microchannels were covered with 2 mL of complete media. Before imaging, a lid was placed on top of the apparatus to prevent evaporation.

### Microscopy

High-resolution imaging was performed using a General Electric (Boston, MA) DeltaVision Elite imaging system mounted on an Olympus (Japan) IX71 stand with a motorized XYZ stage, Ultimate Focus, cage incubator, ultrafast solid-state illumination with excitation/emission filter sets for DAPI, CFP, FITC, GFP, YFP, TRITC, mCherry, and Cy5, critical illumination, Olympus PlanApo N 60X/1.42 NA DIC (oil) and UPlanSApo 60X/1.3 NA DIC (silicone) objectives, Photometrics (Tucson, AZ) CoolSNAP HQ2 camera, SoftWoRx software with constrained iterative deconvolution, and vibration isolation table.

### Cell migration

To perform cell speed and directionality ratio analyses, we used an Excel (Microsoft, Redmond, WA) plugin, DiPer, developed by Gorelik and colleagues and the Fiji plugin, MTrackJ, developed by Erik Meijering for manual tracking^35, 36^. Brightfield imaging was used to confirm that debris were not obstructing the cell.

### Cell stiffness assay

The previously described gel sandwich assay was used with minor modifications^10^. 6-well glass-bottom plates (cat no. P06-1.5H-N; Cellvis, Mountainview, CA) and 18 mm coverslips were activated using 3-aminopropyltrimethoxysilane (Sigma Aldrich) for 5 min and then for 30 min with 0.5% glutaraldehyde (Electron Microscopy Sciences) in PBS. 1 kPa polyacrylamide gels were made using 2 μl of blue-fluorescent beads (200 nm; Thermo Fisher), 18.8 μl of 40% acrylamide solution (cat no. 161-0140; Bio-Rad, Hercules, CA), and 12.5 μl of bis-acrylamide (cat no. 161-0142; Bio-Rad) in 250 μl of PBS. Finally, 2.5 μl of ammonium persulfate (APS; 10%) and 0.5 μl of Tetramethylethylenediamine (TEMED) was added before spreading 9 μl drops onto treated glass under coverslips. After polymerizing for 40 min, the coverslip was lifted in PBS, extensively rinsed, and incubated overnight in PBS. Before each experiment, the gel attached to the coverslip was placed on a 14 mm diameter, 2 cm high PDMS column for applying a slight pressure to the coverslip with its own weight. Then, both gels were incubated for 30 min in complete media before plates were seeded. After placing the bottom gels in plates on the microscope stage, the PDMS column with the top gel was placed over the cells seeded on the bottom gels, confining cells between the two gels. After 1 h of adaptation, the cell height was determined with beads by measuring the distance between gels. The cell diameter was measured using a far-red membrane dye (cat no. C10046; Thermo Fisher). Stiffness was defined as the height (*h*) divided by diameter (*d*).

### Immunofluorescence (IF)

After washing with HEPES buffered saline (HBS), cells in poly-L-lysine (cat no. P4707; Sigma Aldrich) coated 6-well glass-bottom plates (cat no. P06-1.5H-N; Cellvis) were fixed with 4% paraformaldehyde (PFA; Electron Microscopy Sciences) in HBS for 20 min at room temperature. Blocking, permeabilization, antibody incubation, and washing were done in HBS with 1% BSA, 1% fish gelatin, 0.1% Triton X-100, and 5 mM EDTA. A 1:250 dilution of cofilin-1 (cat no. MA5-17275; Thermo Fisher) antibody was incubated with cells overnight at 4°C. After extensive washing, a 1:400 dilution of Alexa Fluor 488-conjugated anti-rabbit secondary antibody (cat no. A-21206; Thermo Fisher) was then incubated with cells for 2 hr at room temperature. Cells were then incubated with a 1:250 dilution of Alexa Fluor 568-conjugated phalloidin (cat no. A12380; Thermo Fisher) and a 1:1000 dilution of DAPI (cat no. D1306; Thermo Fisher). Cells were again extensively washed and then imaged in HBS.

### THP-1 macrophage-like cell differentiation

THP-1 cells were differentiated in complete media with phorbol 12-myristate-13-acetate (PMA; 100 nM) (cat no. 1201/1; Tocris, UK) for 72 hr. After 24 hr of rest in PMA free media, total mRNA was harvested from cells for RT-qPCR analysis.

### Statistics

Sample sizes were determined empirically and based on saturation. As noted in each figure legend, statistical significance was determined by either a two-tailed Student’s t-test or multiple-comparison test post-hoc. Unless otherwise stated, all t-tests or multiple-comparison tests are compared to control, *i.e*., non-targeting, siRNA treated cells. Normality was determined by a D’Agostino & Pearson test in GraphPad Prism 7. * - p ≤ 0.05, ** - p ≤ 0.01, *** - p ≤ 0.001, and **** - p ≤ 0.0001

## Data availability

All data are available upon request to the corresponding author, Dr. Jeremy S. Logue.

## Author Contributions

M.F.U.: Investigation, A.E.D.: Investigation, S.B.L.: Investigation, J.S.L.: Conceptualization, Supervision, Project administration, Visualization, Writing - Original Draft, Funding acquisition

## Acknowledgements

We would like to thank Dr. Nathaniel Cady (SUNY Polytechnic Institute, Albany, NY) for fabrication of the silicon wafers.

## Movies

**Movie 1**. Motile THP-1 monocyte-like cell with LifeAct-EGFP. Microchannels are coated with VCAM-1 (1 μg/mL) and are 3 μm in height, 8 μm in width, and 100 μm in length.

**Movie 2**. Motile THP-1 monocyte-like cell with RLC-EGFP. Microchannels are coated with VCAM-1 (1 μg/mL) and are 3 μm in height, 8 μm in width, and 100 μm in length.

**Movie 3**. Non-motile THP-1 monocyte-like cell with LifeAct-EGFP. Microchannels are coated with VCAM-1 (1 μg/mL) and are 3 μm in height, 8 μm in width, and 100 μm in length.

**Movie 4**. Central focal plane of a non-motile THP-1 monocyte-like cell with LifeAct-EGFP treated with a cofilin-1 siRNA. Microchannels are coated with VCAM-1 (1 μg/mL) and are 3 μm in height, 8 μm in width, and 100 μm in length.

**Movie 5**. Ventral focal plane of a non-motile THP-1 monocyte-like cell with LifeAct-EGFP treated with a cofilin-1 siRNA. Microchannels are coated with VCAM-1 (1 μg/mL) and are 3 μm in height, 8 μm in width, and 100 μm in length.

**Movie 6**. Pseudo colored movie of cofilin-1-EGFP in a motile THP-1 monocyte-like cell. Microchannels are coated with VCAM-1 (1 μg/mL) and are 3 μm in height, 8 μm in width, and 100 μm in length.

